# Structure of the unique tetrameric STENOFOLIA homeodomain bound with DNA

**DOI:** 10.1101/2020.05.08.085126

**Authors:** Prabhat Kumar Pathak, Fei Zhang, Shuxia Peng, Lifang Niu, Juhi Chaturvedi, Million Tadege, Junpeng Deng

## Abstract

Homeobox transcription factors are key regulators of morphogenesis and development in both animals and plants^1^. In plants, the WUSCHEL-related homeobox (WOX) family transcription factors function as central organizers of several developmental programs from embryo patterning to meristematic stem cell maintenance through transcriptional activation and repression^2-4^. The structure of WOX Homeodomain (HD) and the molecular mechanism of its interaction with DNA are unknown. Here, we report the 2.1 Å crystal structure of the STENOFOLIA (STF) HD from *Medicago truncatula* in complex with DNA. STF binds DNA as a novel cooperative tetramer, enclosing nearly entire bound DNA surface. The STF tetramer is partially stabilized by docking of the C-terminal tail from one protomer onto a conserved hydrophobic surface on the head of another in a head-to-tail manner. Helix α3 not only serves a canonical role as a base reader in the major groove, but also provides extensive binding to DNA in the minor groove. Our structural and functional data reveal that STF specifically targets ‘TGA’ sequence and the cooperative tetrameric binding with DNA is key to transcriptional repression in plants. Our data reveal an unprecedented HD:DNA recognition mechanism, representing the first plant HD structure from WOX family of transcription factors.

## Introduction

HD containing transcription factors are one of the most powerful regulators of morphology and differentiation in fungi, animals and plants^5,6^. The *WUCHEL-RELATED HOMEOBOX* (WOX) family is unique to plants^7^, and instructs plant growth and development from a small group of pluripotent cells analogous to the stem cell niche animals. WOX genes play central roles in apical-basal polarity patterning during embryogenesis and maintaining the stem cell niches at various plant meristems during post-embryonic shoot and root growth and lateral organ development such as leaves and flowers^8-14^. WUSCHEL(WUS), the founding member of the WOX family, is a conserved key regulator for shoot apical meristem (SAM) and axillary meristem^3,13,15-17^. WUS paralogs including WOX5 in root apical meristem^4^, WOX4 in procambial/cambial meristem ^18,19^, and WOX1 and WOX3 in leaf marginal meristem^20,21^ perform similar functions. The *Medicago truncatula* WOX1 gene, *STENOFOLIA* (*STF*), and its *Nicotiana sylvestris* ortholog, *LAMINA1* (*LAM1*) regulate leaf blade outgrowth by promoting cell proliferation at the adaxial-abaxial junction through transcriptional repression^22-24^. WUS clade WOX members have a promiscuous ability to substitute for the function of each other if driven by specific promoters as demonstrated by complementing the *lam1* mutant in leaf development^25^ and the *wus* mutant in SAM maintenance^26^, suggesting a conserved mechanism in DNA recognition and transcriptional repression. WUS clade members including WUS and WOX1-WOX7 share a conserved WUS box at the C-terminus, specific to the WUS clade^25,27,28^, and a conserved HD, typical of the whole WOX family^8^. While the HD contacts DNA, the WUS box is essential for recruitment of the TOPLESS (TPL) family transcriptional co-repressors^24,29^. HD has a canonical structure comprised of three-α-helical bundle and is found in a large class of transcription factors ubiquitous in fungi, animals and plants^6^, sharing low sequence identity and variable recognition sequences^30^. A typical HD is about 60 amino acids long, but several types of atypical HD proteins have more or fewer^22,31^, including HD of the WOX family containing 65-70 residues. WUS functions by binding to at least two distinct DNA motifs: The G-box motif, TCACGTGA sequence and the TAAT motif, TTAAT(G/C)(G/C) sequence^9,29,32^. STF can also strongly bind to WUS binding sites and the (GA)/(CT)n elements^33^, indicating conserved motif recognition by WOX HD. Although HDs from other kingdoms of life have been studied structurally^6,34-41^, the structure of WOX HD and its DNA binding mechanism remained elusive. Here, we report the crystal structure of STF HD in complex with dsDNA, representing the first plant HD with a novel tetrameric structure specific to plants.

## Results

### The structure of STF^85-190^:DNA reveals a novel tetrameric HD

Apo STF^86-190^ protein appeared as a monomer in solution (Extended Data Fig. 1). However, it forms a stable complex with DNA in a 1:4 (DNA:protein) stoichiometry (Extended Data Figs. 1, 2). After screening a number of synthetic DNA oligos, we found STF^85-190^ readily crystallized when in complex with a 22-bp DNA promotor sequence. The structure of the complex was determined by single wavelength anomalous dispersion (SAD) using a selenomethione substituted triple mutant (L107M/L110M/L130M) STF protein:DNA complex crystal (see Methods). There are two protein and one DNA molecules in one asymmetric unit of the crystal. The two STF^85-190^ protomers adopt near identical conformation with a root square mean deviation (rmsd) of 1.5Å over 68 equivalent Cα atoms. Together with two additional crystallographically related protein molecules, STF^85-190^ binds DNA as a tetramer (Fig. 1), consistent with the stoichiometry in solution. The structure of STF^85-190^ adopts a canonical HD architecture comprised of a three α-helical bundle core connected by well-ordered loops and a long arm of peptide at the N-terminus, and an additional short helix α4 at the C-terminal tail, with an overall dimension of approximately 42Å x 32Å x 25Å. Helix α3 is significantly longer than other helices and perpendicular to α2, adopting a classical helix turn helix motif.

**Figure 1.**
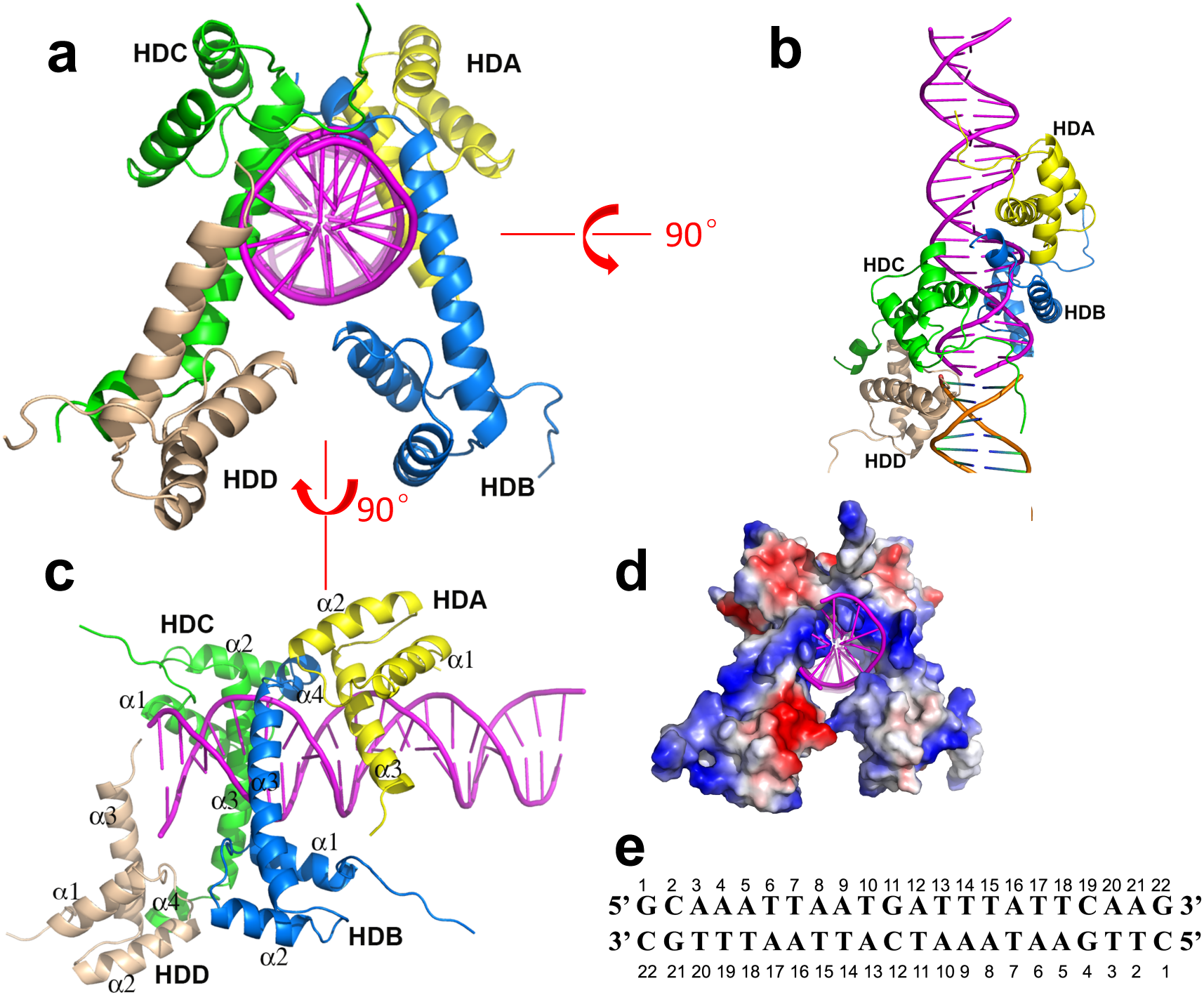
STF^85-190^ binds DNA as a unique tetramer. **a**. Depicted are STF^85-190^ tetramer (HDA in yellow, HDB in blue, HDC in green and HDD in light brown) in complex with a 22-bp DNA (color in magenta). The secondary structures are labeled. **b** and **c** are views from 90° rotations. The secondary DNA forming the pseudo-continuous helix is shown in orange. **d**. The eletropotential surface of STF^85-190^ tetramer is shown. Note, nearly the entire DNA surface is clamped by the protein. **e**. The dsDNA bound sequence from target promoter is shown.

The STF^85-190^ tetramer (HDA, HDB, HDC, HDD) tightly clamps around nearly the entire surface of the DNA spanning three grooves (Fig. 1b, d), burying about 5,435Å^2^ solvent accessible surface (SAS). The tetramer is organized as dimer of dimers (Fig.1) with HDA:HDB dimer packs against HDC:HDD dimer in the DNA major groove. The STF-HD dimers are associated in a head to tail manner involving the short helix α4 at the C-terminus of one protomer docked onto a common hydrophobic surface on the head of the following protomer. This docking pocket is constituted from nonpolar residues located on helix α2 (A120, I123) and α3 (G138, aliphatic side chain of K139, F142 and Y143, Fig. 2c, e). In addition, the tail tip of helix α3 in HDA is associated with helix α1 of HDB in the head via van der Waals interactions, forming a nearly anti-parallel homodimer (Fig. 1c, same in HDC:HDD dimer). The STF-HD tetramer is bridged by helix α4 of HDB, which is sandwiched between HDA and HDC, with one surface involved in contacting HDA (A165, S169 and A170, Fig. 2c), burying about 574 Å^2^ SAS, while the opposite surface involved in docking with HDC (F167 and I171, Fig. 2f) burying about 442 Å^2^ SAS. HDC:HDD interface is same as HDA:HDB (Extended Data Fig. 3).

**Figure 2.**
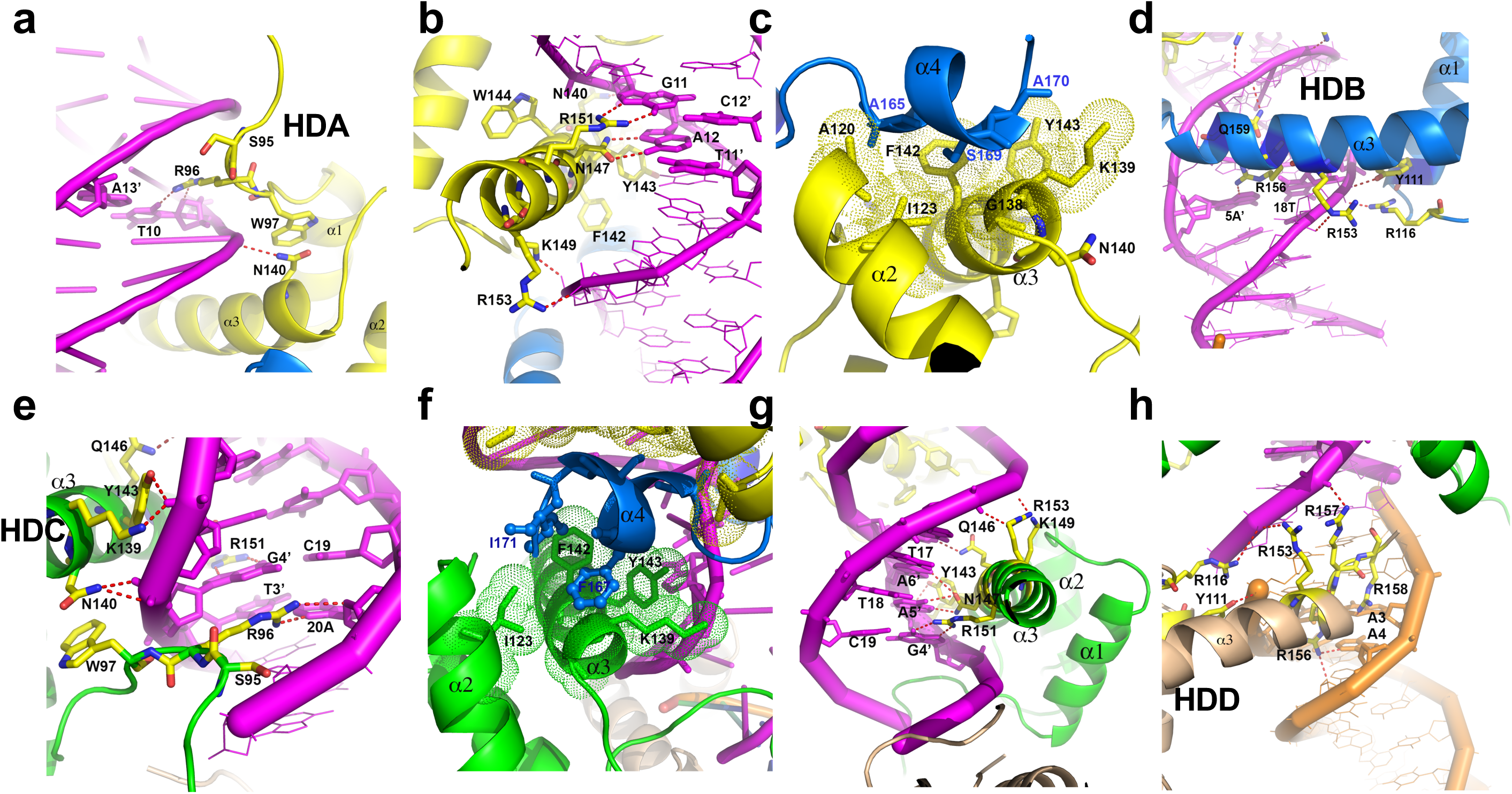
The structure of STF^85-190^:DNA complex. **a**. HDA N-terminal arm showing interaction with DNA minor groove. **b**. HDA helix α3 inserts into DNA major groove and interacts with bases and backbone. **c**. The HDA:HDB dimer interface. The hydrophobic residues lining the docking pocket are shown as sticks, with dotted envelopes indicating the van der Waals radius. **d**. Helix α3 of HDB contacts DNA in the minor groove. **e**. HDC N-terminal arm interacts with DNA minor groove. **f**. HDB:HDC dimer interface. **g**. Helix α3 of HDC contacts DNA in the major groove. **h**. HDD helix α3 contacts DNA in the minor groove. The hydrogen bonds and salt bridges are indicated as red dashed lines. Color scheme is same as in Fig. 1.

### STF^85-190^ recognizes ‘TGA’ DNA sequence

STF^85-190^ tetramer interacts with DNA extensively. HDA and HDC are making contacts with DNA via both minor and major grooves (Fig. 1). Their N-terminal arms are embracing DNA from minor groove, while α3 helices are inserted into the major groove of the DNA. In both HDB and HDD, the α3 helices, the tip of α1 helices, and the α1/α2 loops are contacting DNA via minor groove.

R96 on the N-terminal arm of HDA is inserted into the minor groove of DNA and forms bifurcated hydrogen bonds with O2 and O4’ atoms of base T10 (Fig. 2a). S95 and W97 are bracing the DNA via van der Waals interactions (Fig. 2a). These exquisite interactions contribute to DNA binding affinity, and R96A mutation abolished DNA binding (Fig. 3a). The helix α3 of HDA is sandwiched in the major groove, making extensive interactions with both backbone and bases. Specifically, the N147 side chain is recognizing base A12 through hydrogen bonds with N7 and O6 atoms on the base and the guanidinium head of R151 is hydrogen bonded with N7 and O6 atoms of base G11 (Fig. 2b). Therefore, N147 and R151 together could serve as a molecular probe for recognizing the ‘TGA’ DNA fingerprint. In addition to these base specific interactions, helix α3 are contacting the backbone of DNA via hydrogen bonds and salt bridges (N140, K149 and R153), as well as hydrophobic interactions with the DNA bases (F142, Y143) and backbone (K139, W144).

**Figure 3.**
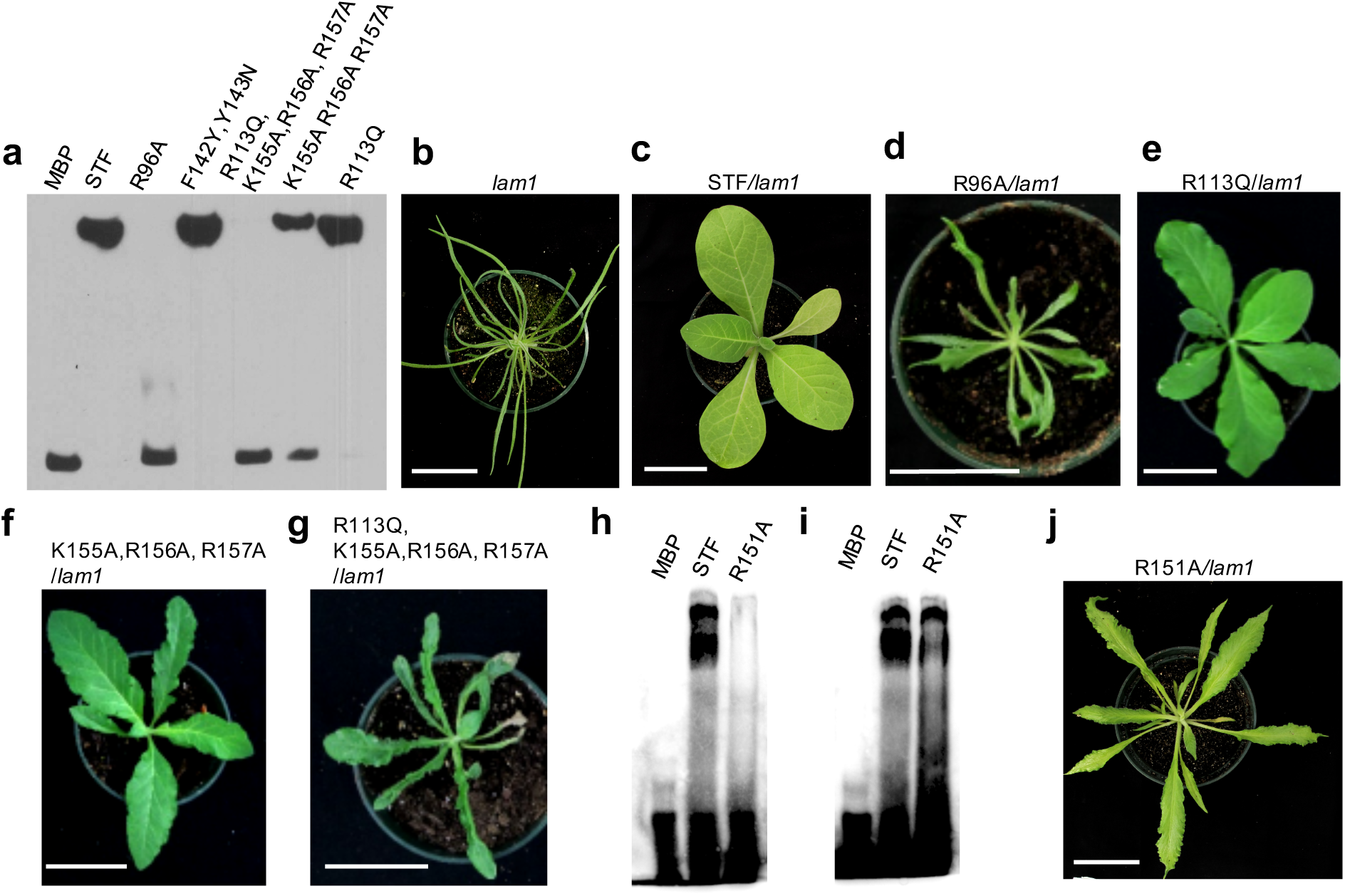
Key residues of STF-HD for DNA binding and *in vivo* function. **a**. EMSA showing that mutations in STF-HD affect the DNA binding ability *in vitro*. **b-g**. Phenotypes of *lam1* mutant (**b**) plants complemented with wild type *STF:STF* (**c**), or mutants *STF:STF-R96A* (**d**) *STF:STF-R113Q* (**e**), STF:STF-K155AR156AR157A (**f**), STF:STF-R113QK155AR156AR157A (**g**). **h**. EMSA showing that R151A mutation nearly abolished STF’s binding to the “TGA” sequence (GCAAATCTATGATCTATTCAAG). **i**. EMSA showing that R151A mutation only reduced STF’s binding to “TAAT” sequence (GCAAATTAATTATTTATTAAAG). **j**. Phenotype of *lam1* mutant plant complemented with *STF:STF-R151A*.

With its helix α4 tethered on the head of HDA (Fig. 2d), HDB is contacting DNA mainly in the minor groove via basic and polar residues from helix α3 (Q159, R153 and R156), α1/α2 loop (R116) and helix α1 (Y111). Except R156 that is hydrogen bonded with N3 atom on base A5’ (reverse strand) in the minor grove, all others are making DNA backbone contacts via hydrogen bonding and salt bridges (Fig. 2d). The insertion of helix α4 of HDB into the major groove causes a slight wideness of the groove and a minor kink on the DNA backbone.

The N-terminal arm of HDC interacts with the minor groove of DNA in a similar way as observed in HDA. R96 forming hydrogen bonds with base A20, while flanking S95 and W97 embrace the DNA via van der Waals interactions (Fig. 2e). With its head tethered with the tail of HDB (Fig. 2f), helix α3 of HDC is sandwiched in the major groove of the DNA, forming extensive interactions with DNA similar to HDA (Fig. 2g). N147 and R151 are again serving as a base reader, recognizing A5’/G4’ step. NE2 of Q146 is hydrogen bonded with O4 atom on base T7’. This interaction may not be base specific since OE1 of Q146 could be hydrogen bonded with N4 of a cytosine base. K139, F142 and Y143 are embracing the backbone and bases through van der Waals interactions. The charged and polar heads of K149, Y143 and R153 on helix α3 are also binding DNA backbone through hydrogen bonding and salt bridges (Figs. 2f, 2g).

While the head of HDD is tethered with the tail of HDC (Extended Data Fig. 3), the tip of the helix α3 of HDD is inserted into the following minor groove at the junction between two pseudo-continuous DNA molecules (Fig. 2h). HDD mainly interacts with the backbone of the DNA via a cluster of basic residues (R116 on α1/α2 loop, R153 on helix α3, and R157 and R158 on α3/α4 loop), with the addition of hydrogen bond contributed from Y111 on helix α1 (Fig. 2h). In addition, R156 on helix α3 is forming bifurcated hydrogen bond with N3 of base A4 and O4’ of base A5 on the subsequent DNA molecule. These interactions may contribute to DNA binding affinity.

### Structure based mutagenesis: key residues for DNA recognition and tetramer organization

Based on the current complex structure, we carried out mutagenesis to identify key residues of STF^85-190^ that are essential for DNA binding and STF function. We found R96A mutation nearly abolished DNA binding and *lam1* mutant complementation (Fig. 3a-d). Mutation of R113Q, on the other hand, did not affect the DNA binding, which is consistent to the observation that R113 not significantly involved in DNA binding (Fig. 3a, e). Triple alanine substitutions of the positive charge cluster on helix α3, KRR/AAA (155-157) reduced the binding, and the combination of KRR/AAA and R113Q mutations greatly reduced the DNA binding and STF’s ability to rescue the *lam1* mutant (Fig. 3a, f, g).

The STF protein can bind DNA sequences with either “TGA” or “TAAT” specificity (Extended Data Figs. 4, 5). In the STF^85-190^-DNA crystal structure, R151 is specific to bind the guanine of the second base pair of the ‘TGA’ sequence, although the “TAAT” box is also present in the 22bp DNA (Fig. 1e). To identify the determinant motif, we tested STF binding with either “TGA” or “TAAT” sequences. The R151A mutation abolished binding to the “TGA” containing DNA, while retained binding to “TAAT” at a reduced level (Fig. 3h, i). The R151A mutant significantly lost the ability to complement the *lam1* mutant (Fig. 3j), and disrupted tetrameric binding to DNA in EMSA (Extended Data Fig. 6), indicating that R151-mediated STF HD binding to the TGA motif is crucial for STF function. To evaluate the significance of STF tetramer in DNA binding and STF function *in planta*, we carried out mutagenesis at the tetramer interface based on the structure. Specifically, we substituted the two highly conserved aromatic residues at the docking pocket, F142 and Y143, which not only provide a platform for accepting the helix α4 of the neighboring protomer, but also embrace DNA bases via van der Waals interactions to provide affinity. We found that the F142Y/Y143N double mutation abolished the cooperative tetrameric binding to DNA in EMSA-based agarose gel shift assay (Extended Data Fig. 7), and reduced the STF repressive activity (Extended Data Fig. 8a, b), leading to reduced *lam1* complementation (Extended Data Fig 8c, d). Similarly, N147I mutation abolished its binding to DNA and *lam1* complementation (^24^ and Extended Data Fig 8d), suggesting essential roles of R96, F142, Y143, N147 and R151 residues for the STF tetrameric structure and function.

## Discussion

WOX family proteins are plant specific transcription factors that play central roles as master regulators of zygotic and embryonic patterning, stem-cell maintenance and lateral organ development^3,4,20-22,42^. However, the structure and mechanism of plant HD recognition of DNA remains elusive. Here we report the first WOX-HD crystal structure, in which the STF helix α3 not only recognizes DNA from major groove as the single base “recognition” helix seen in all known HDs^30^, but also provides extensive interactions with DNA from minor groove (Fig. 2). STF^85-190^ probes the DNA bases in the minor groove with R96 on its N-terminal arm, and in the major groove with two residues, N147 and R151 on helix α3 similar to other HDs. Although the STF-HD could recognize both the ‘TAAT’ and TGA(X)_2-5_TCA motifs (Fig. 1e), STF^85-190^ strongly binds only to the ‘TGA’ motif in the current structure. This is consistent with reports indicating that the binding affinity of the TGA(X)_2-5_TCA is 20-fold higher than the ‘TAAT’ motif^29^, and STF preferentially targets GA/TC sequences^33^. This difference in binding affinity could therefore offer WOX-HD to bind to specific targets distinct from other HD proteins.

While all other HD:DNA structures solved to date involve either monomer or dimer HDs^6,43^, STF:DNA complex revealed an unprecedented tetrameric configuration. The tetramer clamps to the DNA over nearly the entire surface of the bound DNA region, in contrast to just a portion of bound DNA surface observed in other structures. This unique STF homo-tetramer resulted from cooperative binding to substrate DNA, and is stabilized by the bridging helix α4 as a C-terminal extension to the canonical HD core. The STF-HD tetramer is organized as a dimer of dimers with each dimer displaying a unique head to tail antiparallel unit (Fig. 1c). Although head to tail type of association was observed in yeast MATa1 and MATα2 HD heterodimer bound with DNA^39^, in contrast to the helix α4 of MATα2 bound on the surface of helix α1 and α2 in MATa1, the helix α4 of one STF HD is docked on helix α2 and α3 at the head of the other, forming a unique tetramer. In addition, the nearly antiparallel dimer association of STF HD in a head to tail manner is unprecedented.

The WOX family has been phylogenetically divided into WUS/ modern clade, WOX9/intermediate clade, and WOX13/ancient clade with transcriptional repression activity in the WUS and activation activity in the WOX9 and WOX13 clades^25,26,44,45^. While the STF-HD tetramer is typical of the WUS clade, it is yet to be shown if DNA binding as cooperative tetramer is a feature of the entire WOX family. The STF G138 and K139 residues replacement with D138 and A139 in WOX9 could potentially compromise its interactions with DNA backbone, weakening the tetramer association. In addition, WOX13/ancient clade HDs have a F142Y and Y143N substitution (^44^ and Extended Data Fig. 9), which could drastically weaken both its DNA binding and tetramer association (Extended Data Fig. 7). Thus, it will be interesting to see if the WOX-HD offers explanation to the evolutionary dynamic nature of WOX family proteins, in addition to the diagnostic WUS box that recruits TPL for transcriptional repression. TPL forms a tetramer for its co-repressor function and the oligomeric states of repressors could dramatically alter the TPL binding affinity^46^. This suggests that the STF-HD tetramer could enhance its association with TPL by multivalent interactions, conferring preferential advantage to WUS clade WOX proteins. Our data uncovers a novel HD:DNA recognition mechanism and provides mechanistic insight into the function of dynamic WOX genes and their contribution to the complex morphology and developmental evolution of higher plants.

## Methods

### Protein purification and crystallization

The coding sequence of *Medicago truncatula* STENOFOLIA_85-190 residues was amplified by PCR and inserted into a modified PET vector as a MBP fusion with a N-terminal 6XHis-tag that is cleavable by Tabaco etch virus protease (TEV). The recombinant protein was expressed in *E.coli* and purified by Ni-NTA as previously described^47^. Briefly, STF^85-190^ protein was first purified from soluble cell lysate using Ni-NTA affinity column. The eluted protein was subsequently subjected to TEV protease cleavage and was collected as flow through of a second subtracting Ni-NTA column. The protein was further purified by size exclusion chromatography and cation exchange purification to homogeneity. Mutant STF proteins were purified same as WT. The purified proteins were concentrated to 20-25 mg/ml in buffer 20 mM Tris-HCl, pH 7.4, 125 mM NaCl and 5 mM TCEP [Tris (2-carboxyethyl)phosphate], flash frozen and stored at - 80 °C until usage^48^. STF^85-190^ L107M/L110M/L130 M triple mutant was cloned using the PCR-based site-directed mutagenesis method. The selenomethionine (SeMet) substituted proteins were expressed in *E.coli* BL21(DE3) with SeMet supplemented in M9 medium, and purified using the same procedures as described above.

The 22-bp synthetic oligonucleotides containing the sequence 5’-GCAAATTAATGATTTATTCAAG-3’ and its complementary oligonucleotide 5’-CTTGAATAAATCATTAATTTGC-3’ were annealed in buffer containing 50 mM HEPES, pH7.2, 50 mM NaCl, 5 mM MgCl_2_, with a temperature gradient from 95°C to 23°C in 2 hours. STF^85-190^ was mixed with the 22-bp DNA at 4:1 molar ratio before crystallization trials. The complex crystals for both WT and SeMet substituted triple mutant crystals were both obtained from a condition containing 0.15 M sodium chloride 28% v/v PEG Smear Medium at 20 °C. 20% glycerol was added to the mother liquid as cryoprotectant.

### Structural determinations

All data were collected at the beamline 19-ID at the Advanced Photon Source (APS), Argonne National Laboratory. Our attempts of using molecular replacement method to solve the native data set using canonical HD domain structures as templates failed. Selenomethionine (SeMet) substitution of WT protein could not yield usable anomalous signal to solve the structure due to disorder of the single M160 present in the protein. Based on the homology modeling with HD domains, we made a triple mutant of STF by substituting three buried leucine residues with methionines (L107M/L110M/L130M). The structure of STF^85-190^:DNA complex was solved by Single-Wavelength-Anomalous-Dispersion method using program HKL3000^49^, with data collected from a single SeMet substituted triple mutant protein crystal. 70% of all protein residues were constructed from the experimental phases obtained from the SeMet crystal data using the program Autobuild in PHENIX^50^. The remaining residues and the 22-bp DNA were built manually using COOT^51^. This model was used to solve the native structure at higher resolution by molecular replacement method using program Phaser^52^. PHENIX program^50^ was used for the refinement, and COOT^51^ was used for the iterative manual model building. Translation, libration and screw-rotation displacement (TLS) groups used in the refinement were defined by the TLMSD server^53^. The final *R*_work_ and *R*_free_ for the refined model were 19.4% and 25.0%, respectively. The current model is of good geometry and refinement statistics (Extended Data Table 1). All molecular graphic figures were generated with PYMOL^54^.

### EMSA for protein:DNA binding

Purified STF^85-190^ proteins were tested for DNA binding on agarose gel based EMSA. 6-FAM labeled DNA oligos (*IDTdna*) was mixed with purified proteins at various molar ratios and incubated on ice for 60 minutes before electrophoresis on 1% agarose gel in TAE buffer for 60 min at 90V at 4°C. The gel was subsequently analyzed on a Biorad ChemiDoc fluorescence imager with 497 nm Ex and 520 nm Em wavelengths.

STF^85-190^ proteins binding with DNA were also analyzed by native polyacrylamide gel based EMSA as previously described^55^. Briefly, oligos were synthesized with the 3’ Biotin CPG modification. Oligos were annealed and incubated with His-MBP, His-MBP-STF or His-MBP-STF mutant fusion proteins using the Light Shift Chemiluminescent EMSA Kit (*Pierce*) at room temperature for 30 min. The binding reaction was: 1xbinding buffer, 2.5% glycerol, 5 ng/ µL Poly (dI.dC), 0.05% NP-40, 50 mM KCl, 0.05 µg/µL purified protein, 5 fmol/µL annealed oligos. Gel electrophoresis was performed on a 5% native polyacrylamide gel. After blotting on a positively charged nylon membrane, the DNA was cross-linked using a transilluminator at standard condition. The biotin-labeled DNA was then detected by using the Chemiluminescent Nucleic Acid Detection Module Kit (*Pierce*).

### Plant Materials and Growth Conditions

The *Nicotiana sylvestris* (*N. sylvestris*) wild type *and lam1* mutant were used in this research. Plants were grown in a controlled greenhouse with 24°C/16-h (day) and 20°C/8-h (night) photoperiods, 60%-70% relative humidity, and 150 µmol m^-2^ s^-1^ light intensity.

### Plasmid Construction and Plant Transformation

All *lam1* complementation assays were performed by using the pSTF-pMDC32 Gateway vector as described^25^. The mutations in STF were introduced using appropriate mutagenic primers and were confirmed by sequencing. STF and the mutated forms were cloned to pDONR207 vector and then ligated to the pSTF-pMDC32 destination vector by LR reaction (*Invitrogen*). Constructs were introduced into *Agrobacterium tumefaciens* strain GV2260 for *N. sylvestris* transformation. Leaf blades from 2-month old *lam1* mutant were used for the transformation. The transformation was performed as previously described^22^. The complementation strength was evaluated by the leaf length/width ratio of the largest leaf in each independent transgenic lines (Extended Data. Fig. 8). At least 10 independent lines were analyzed for each construct.

## Acknowledgements

We gratefully acknowledge the staff of beam-line 19ID at the Advanced Photon Source for their support. This project was supported by the Oklahoma Agricultural Experiment Station at Oklahoma State University under project OKL03060 to J.D and by the National Science Foundation under grant IOS-1354422 to M.T. The authors declare no competing interest.

## Author Contributions

J.D. and M.T. designed research; P.P. and S.P. determined the STF^85-190^:DNA structure. F.Z. and L.N. carried out mutagenesis, DNA binding assays and transgenic plant studies; P.P. and J.C. carried out protein purification and DNA binding studies; S.P., F.Z., M.T. and J.D. analyzed data and wrote the paper.

## Author Information

Atomic coordinates and structure factors have been deposited with the Protein Data Bank, www.rcsb.org, with accession codes 6WIG. Reprints and permissions information is available at www.nature.com/reprints. The authors declare no competing financial interests. Correspondence should be addressed to J.D.

## Extended Data

**Extended Data Figure 1.**
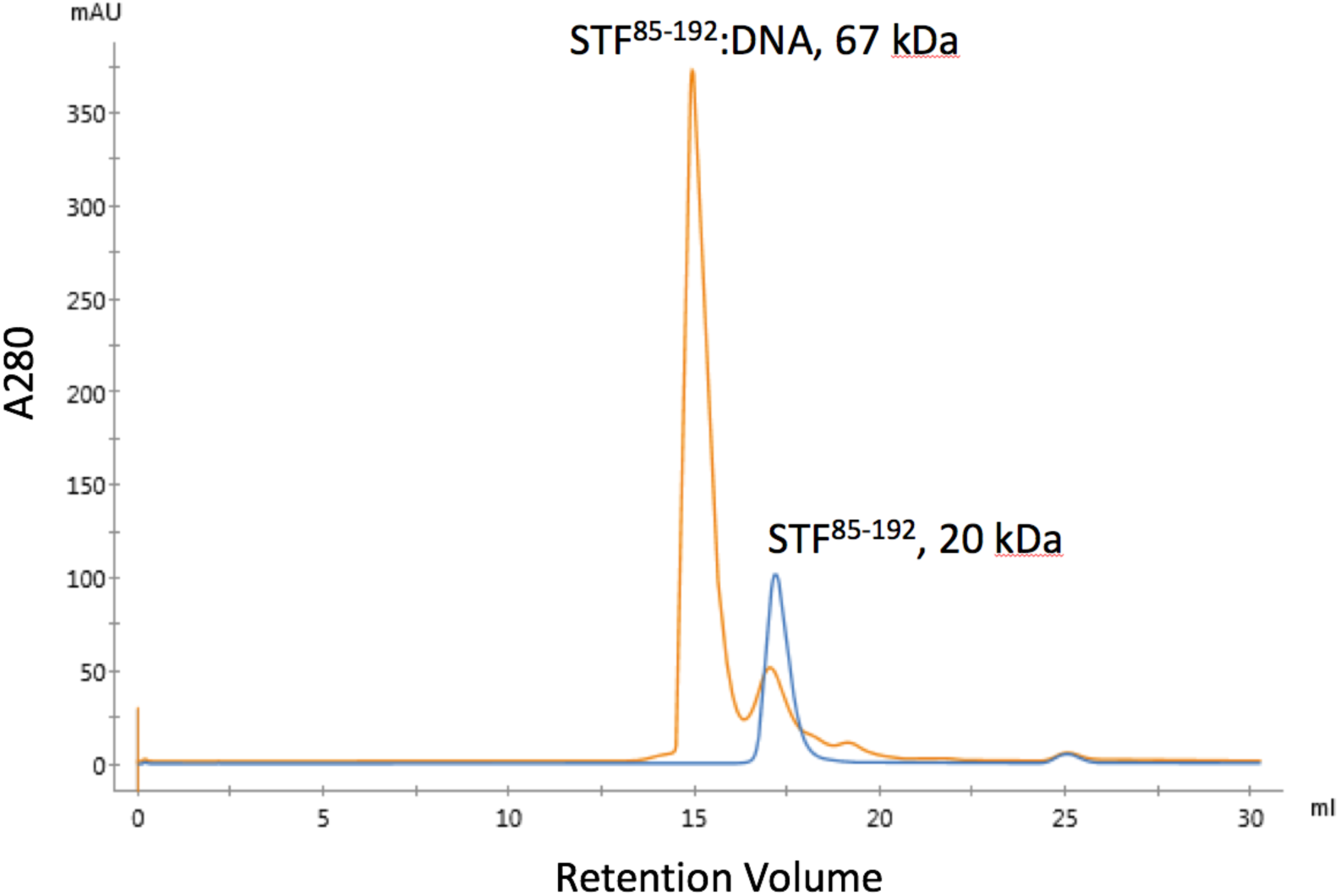
Gel filtration analysis of STF^85-190^ and its complex with DNA. Chromatographs of apo protein (blue) and the DNA complex (orange) from a superdex s200 column are shown. The estimated MW of apo protein from the retention volume is about 20 kDa, which is larger than the theoretic MW of 12.5 kDa of STF ^85-190^ due to its elongated shape. This data is in agreement to the STF^85-190^ monomer with calculated apparent MW of 22.5kDa in solution (www.fluidic.com) based on its calculated hydrodynamic radius of 23.4Å using the current crystal structure^56^. The peak faction collected from the complex was tested for A280/A260 with 1.61 value, suggesting 1 DNA:4 protein in the complex.

**Extended Data Figure 2.**
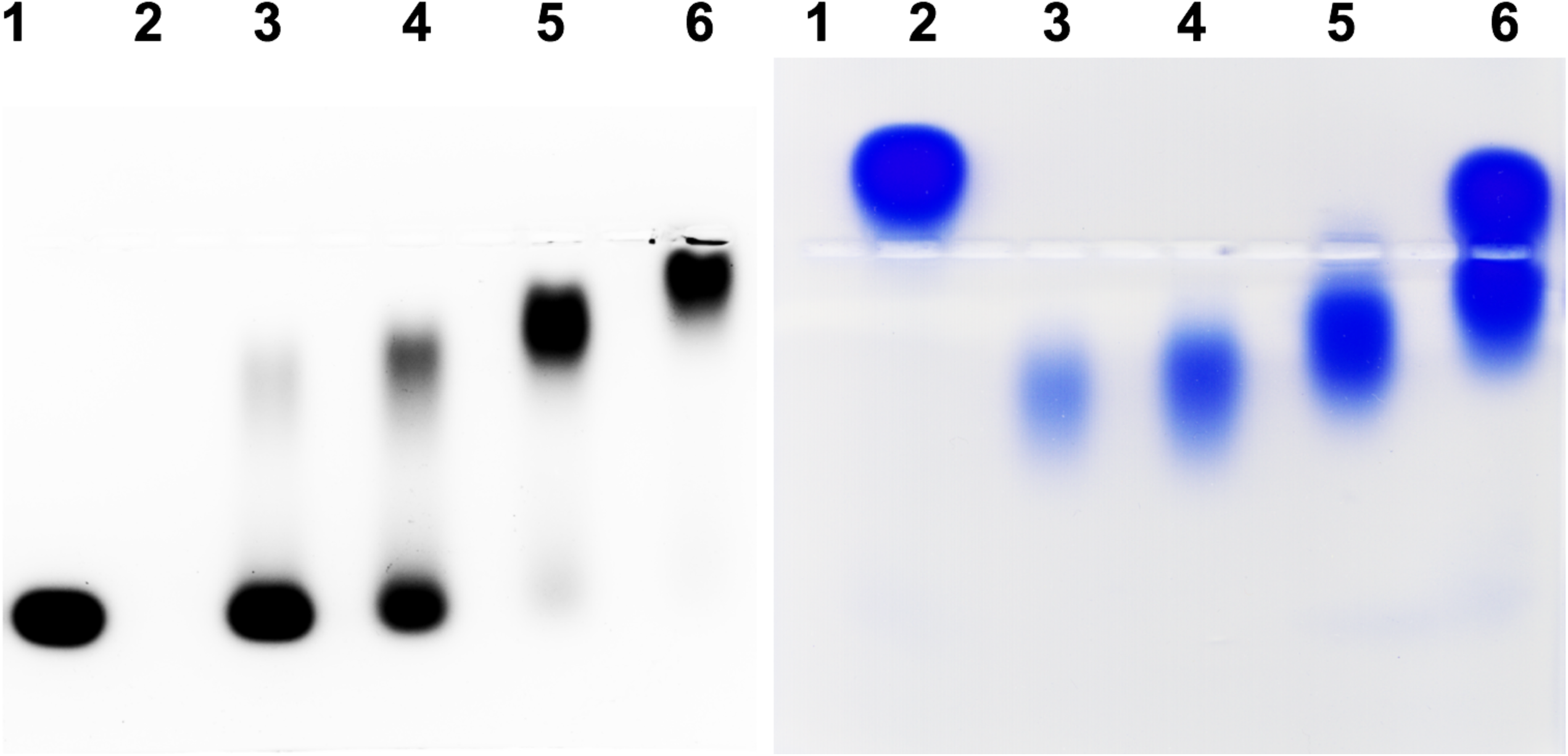
EMSA analysis of STF^85-190^ binding with 6-FAM labeled DNA. Left, the fluorescence signal from the DNA is captured. Right, the same gel is stained with coomassie blue to show the protein. Lane 1, 6-FAM labeled DNA; 2, apo protein; 3-6, DNA:protein mixed at molar ratios of 1:1, 1:2, 1:4 and 1:8 respectively. Note that STF^85-190^ forms stable complex with DNA at 1:4 molar ratio, lane 5.

**Extended Data Figure 3.**
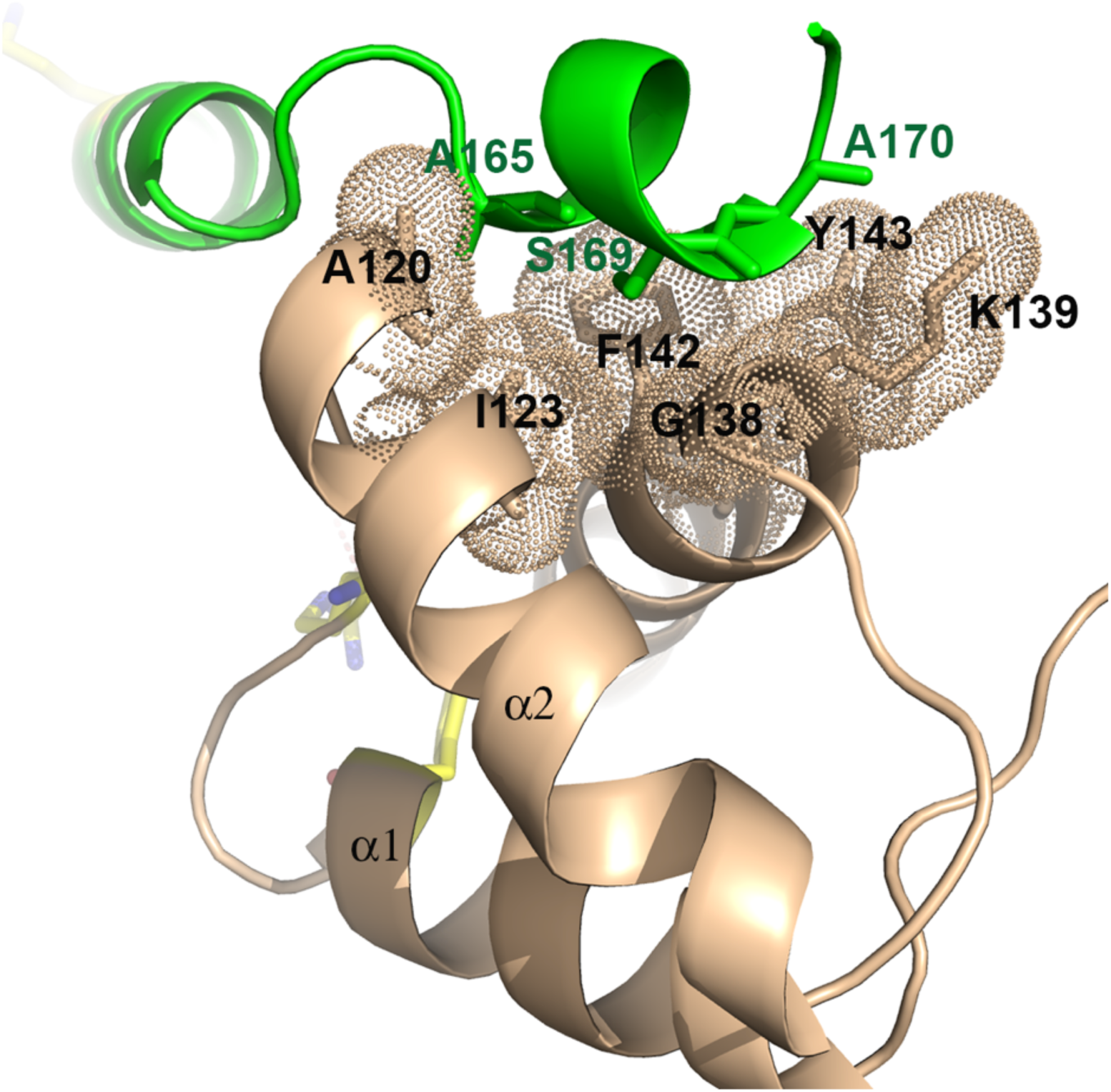
The HDC:HDD dimer interface. The helix α4 of HDC is shown in green and the residues on the hydrophobic surface of HDD (colored in light brown) are shown as sticks with dotted envelopes indicating van der Waals radius. This interface is same as observed in HDA:HDB.

**Extended Data Figure 4.**
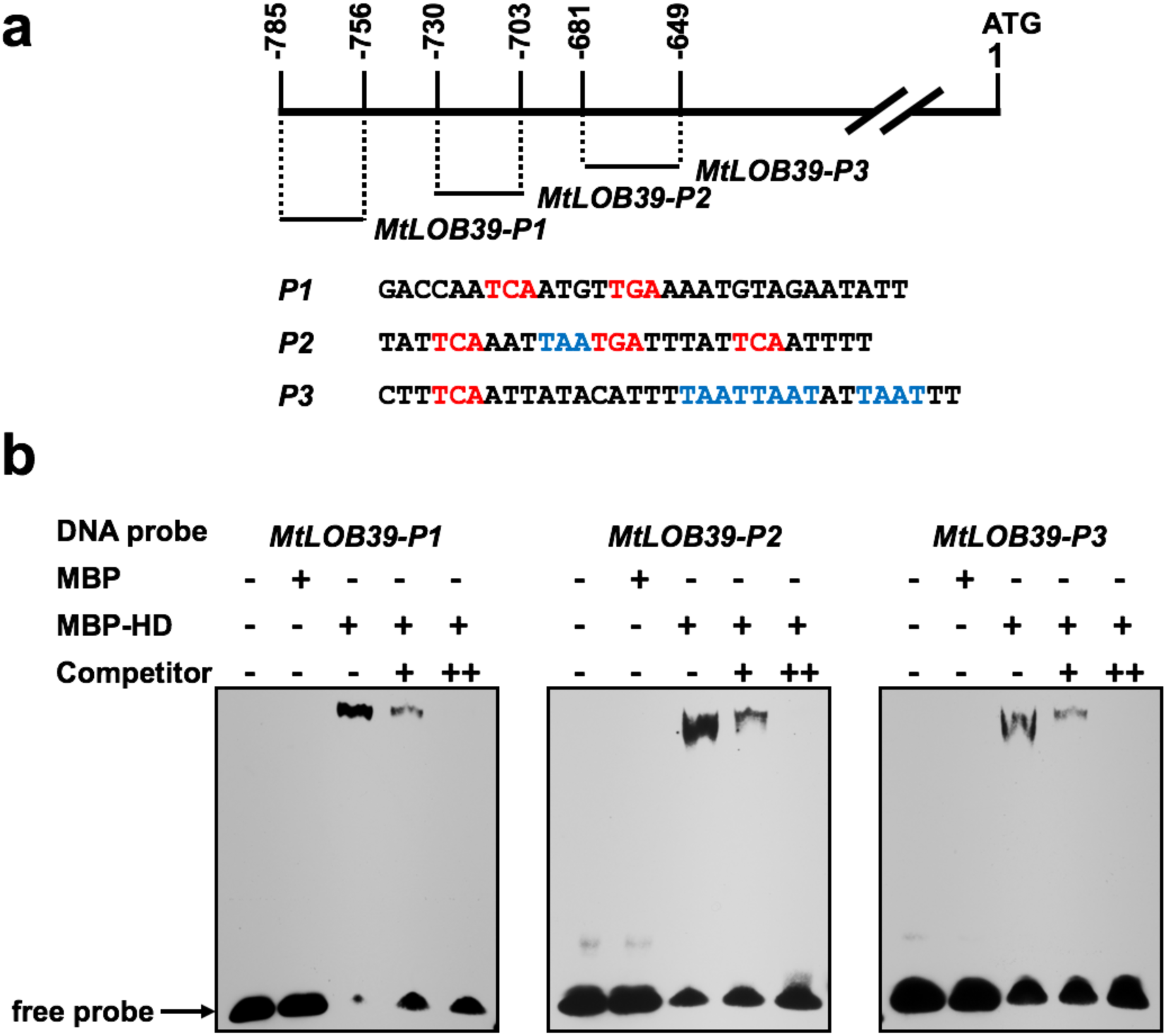
STF-HD specifically interacts with *MtLOB39* promoter regions. **a**. Diagram of *MtLOB39* promoter regions and the sequences containing “TGA” (red) and/or “TAAT” (blue) core sequences. **b**. EMSA showing that MBP-STF-HD specifically binds to all the P1, P2, P3 regions of the *MtLOB39* promoter. Fifty-fold excess unlabeled same oligos were used as competitors to show specific binding.

**Extended Data Figure 5.**
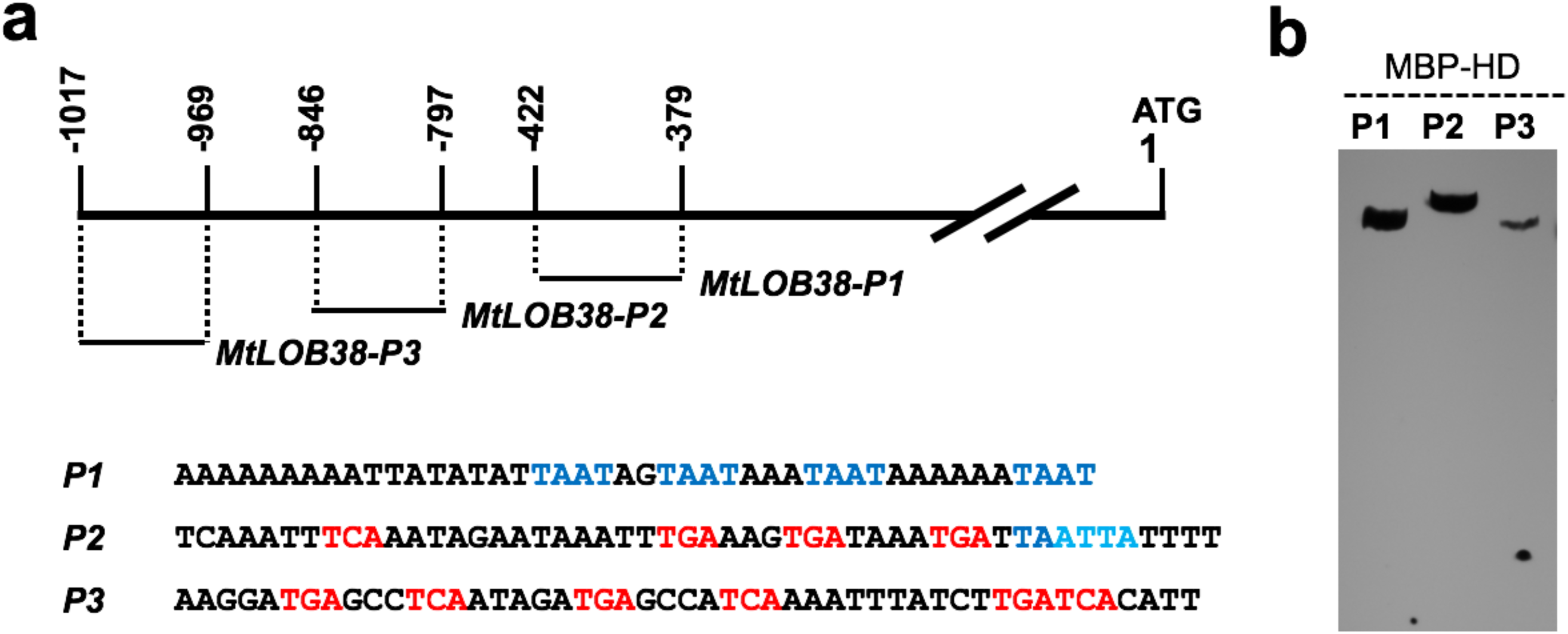
STF-HD specifically interacts with *MtLOB38* promoter regions. **a**. Diagram of *MtLOB38* promoter regions and the sequences containing “TGA” (red) and/or “TAAT” (blue) core sequences. **b**. EMSA showing that MBP-STF-HD binds to all the P1, P2, P3 regions of the *MtLOB38* promoter.

**Extended Data Figure 6.**
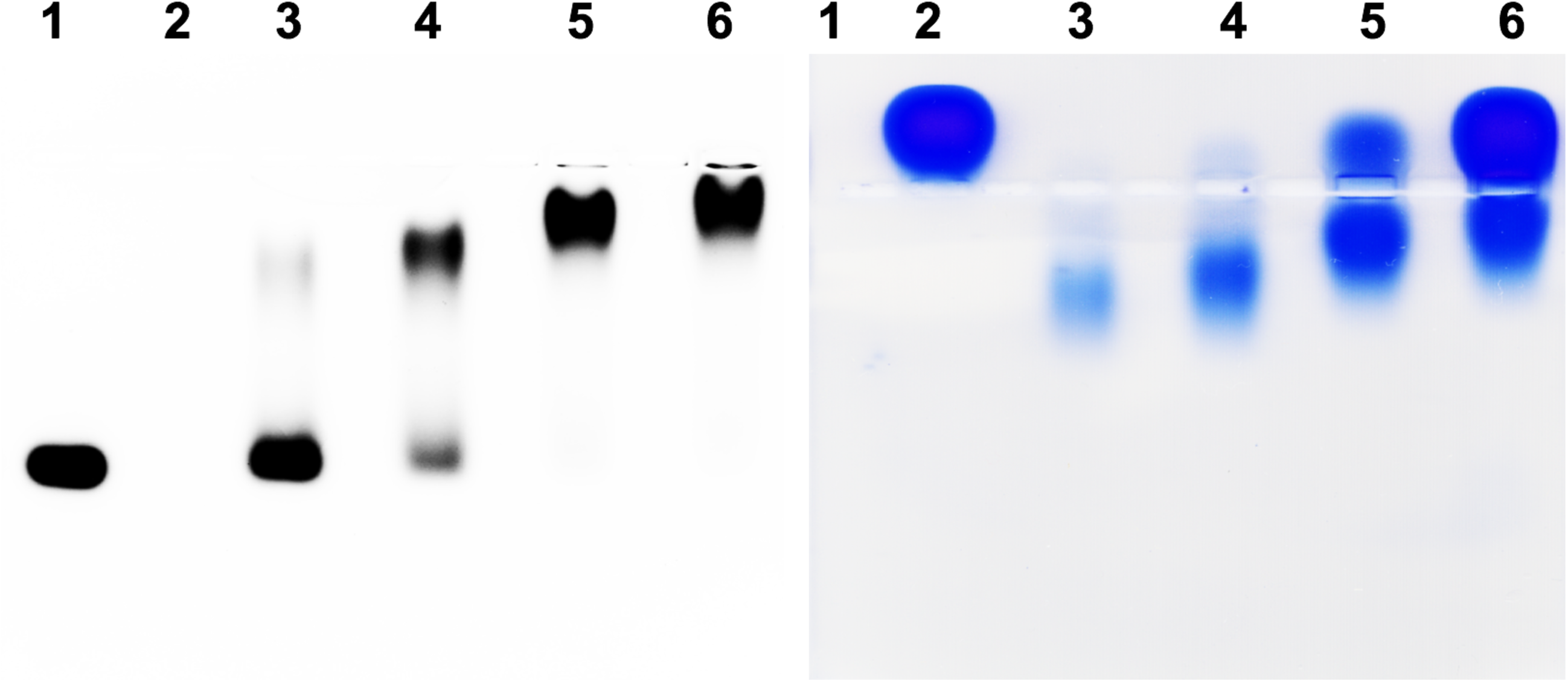
EMSA analysis of STF^85-190^ R151A mutant binding with 6-FAM labeled DNA. Left, the fluorescence signal from the DNA is captured. Right, the same gel is stained with coomassie blue to show the protein. Lane 1, 6-FAM labeled DNA; 2, apo protein; 3-6, DNA:protein mixed at molar ratios of 1:1, 1:2, 1:4 and 1:8 respectively. Note that STF^85-190^ shifted most of the DNA at 1:2 molar ratio (lane 4), indicating the dissociation of the tetramer.

**Extended Data Figure 7.**
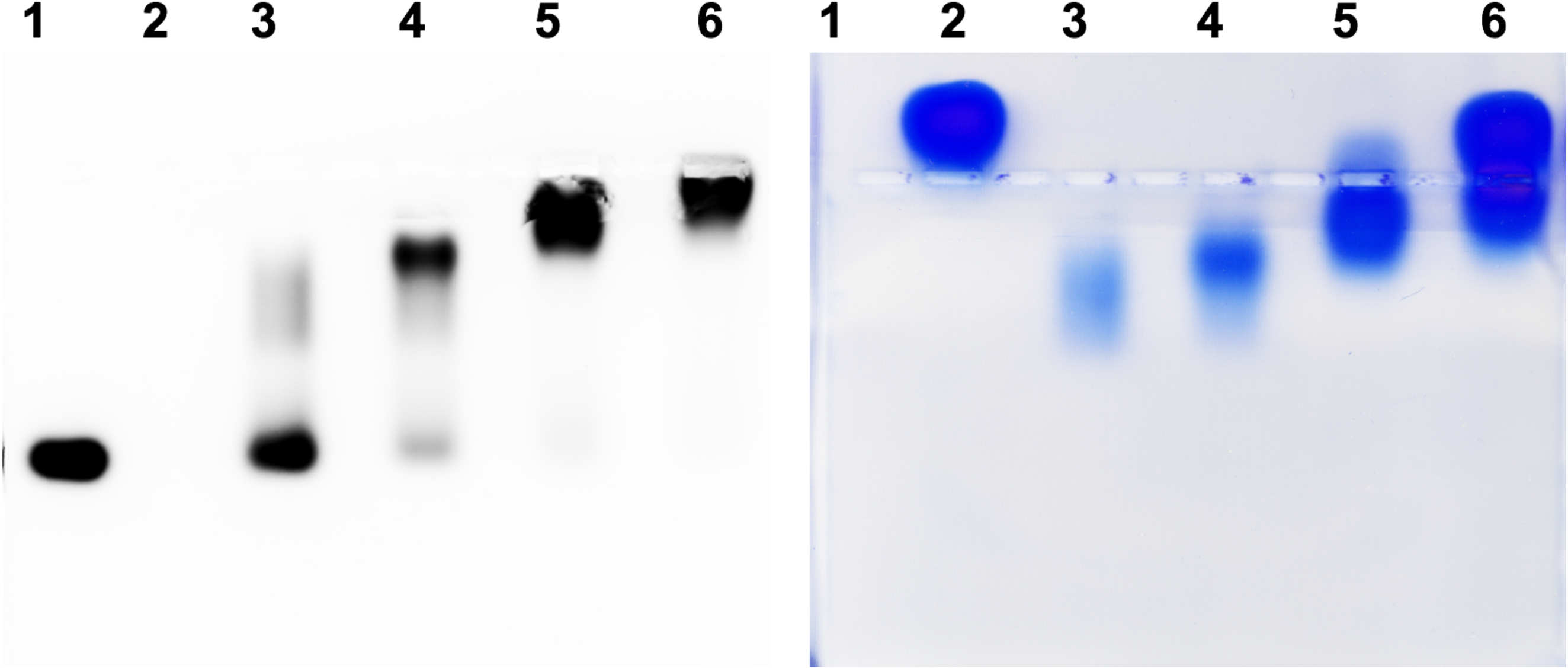
EMSA analysis of STF^85-190^ F142Y/Y143N mutant binding with 6-FAM labeled DNA. Left, the fluorescence signal from the DNA is captured. Right, the same gel is stained with coomassie blue to show the protein. Lane 1, 6-FAM labeled DNA; 2, apo protein; 3-6, DNA:protein mixed at molar ratios of 1:1, 1:2, 1:4 and 1:8 respectively. Note that STF^85-190^ shifted most of the DNA at 1:2 ratio (lane 4), indicating the dissociation of the tetramer.

**Extended Data Figure 8.**
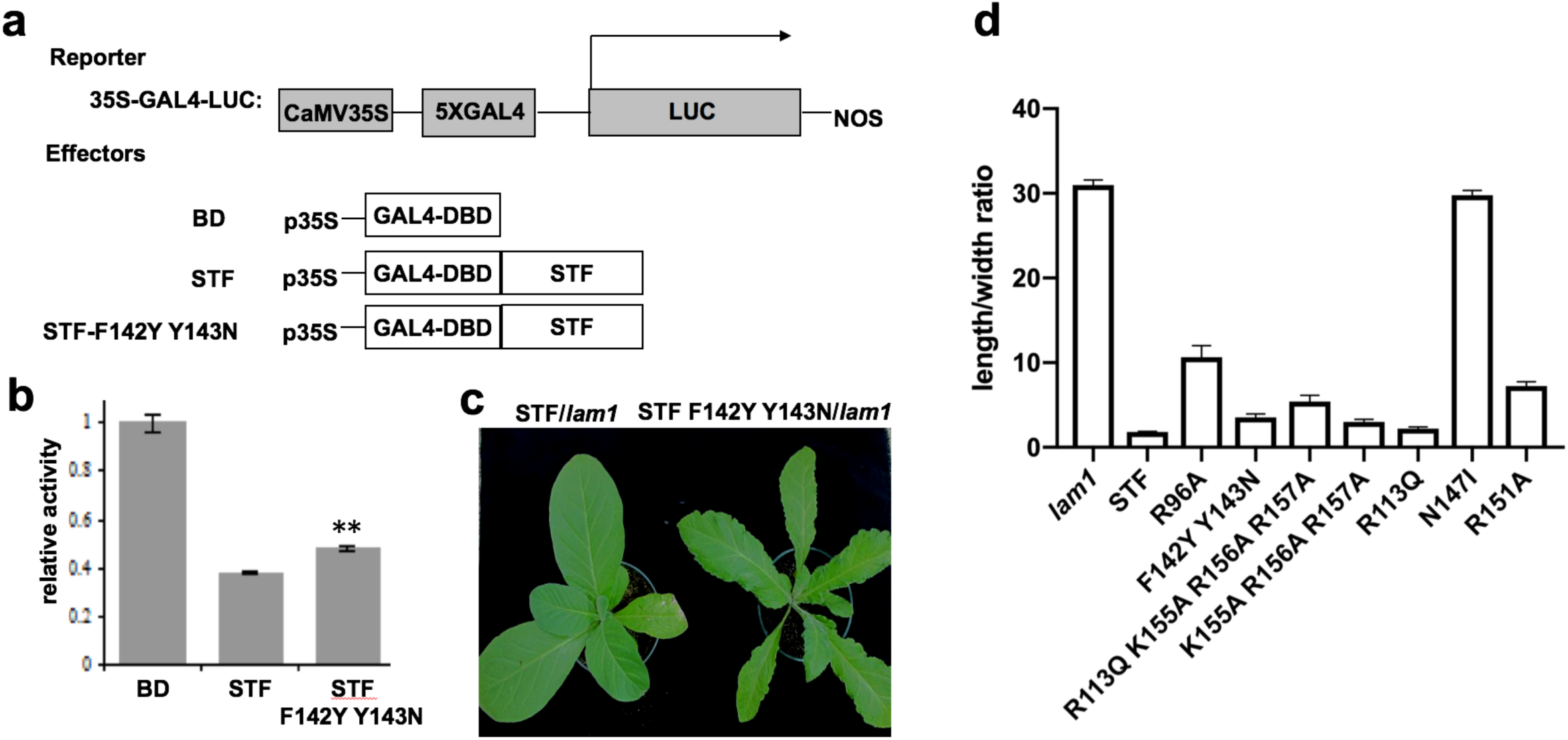
STF F142Y/Y143N mutant has reduced repressive activity and compromised biological function. **a**. Diagram of reporter and effector constructs used in dual luciferase assays. **b**. Relative activity of STF or STF F142Y/Y143N mutant on the reporter. STF F142Y/Y143N mutant showed significantly reduced repressive activity compared to STF. Standard errors were calculated from the means of three biological replicates, each runs in triplicate. **, P<0.01 (t-test). **c**. STF F142Y/Y143N showed reduced activity in complementing the *lam1* narrow leaf phenotype. **d.** Complementation of *lam1* mutant by STF with or without mutations in the homeodomain. The leaf length/width ratio was calculated from the largest leaves of each plant, at least 10 independent lines of each construct were measured.

**Extended Data Figure 9.**
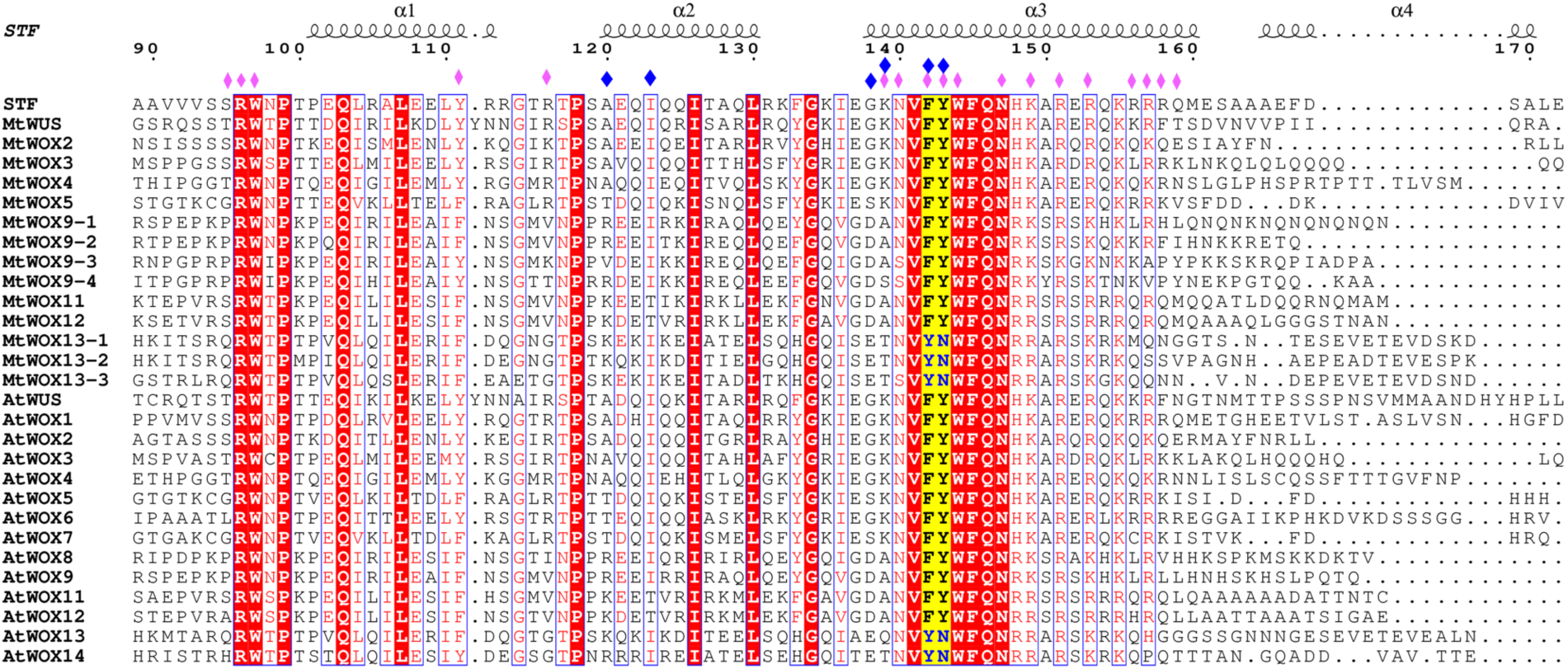
Structure based sequence alignment of selected WOX HD domains. Structure based sequence alignment of various STF HD orthologs was created using the crystal structures of STF^85-190^ as the template. Lettering and numbering above the alignment correspond to STF^85-190^ topology and numbering scheme. Sequence alignment was performed with SSM server ^57^, and the figure was created with ESPript ^58^. Residues involved in DNA binding are indicated with purple diamonds and residues that constitute the docking platform at the dimer interface are indicated with blue diamonds.

---

**Extended Data Table I.** Crystallographic data and statistics.

| Data collection | SeMet_STF_M_Peak | Native_STF |
| --- | --- | --- |
| Beamline | 19-ID, APS | 19-ID, APS |
| Wavelength, Å | 0.97918 | 0.97935 |
| Space group | P2<sub>1</sub> | P2<sub>1</sub> |
| Cell parameters a, b, c, Å | 46.3, 49.0, 70.0 | 48.1, 49.5, 69.8 |
|  | β=105.9° | β=106.5° |
| Resolution, Å | 50.00–2.50 (2.59–2.50) | 50.00–2.10 (2.18–2.10) |
| Total reflections | 55,241 | 142,294 |
| Unique reflections | 10,086 (794) | 17,554 (1,431) |
| Redundancy | 5.5 (3.6) | 8.1 (5.8) |
| Completeness, % | 94.1 (74.0) | 95.2 (77.7) |
| I/σ | 14.2 (2.5) | 23.5 (1.9) |
| Rsym, % | 13.0 (48.6) | 8.0 (66.2) |
| CC1/2, % | 95.4 (81.6) | (81.4) |
| Refinement statistics |  |  |
| Resolution range used, Å |  | 46.1–2.1 |
| No. reflections used |  | 17,488 |
| Rwork/Rfree, % |  | 19.4/25.0 |
| Rmsd bond lengths, Å |  | 0.010 |
| Rmsd bond angles, ° |  | 1.214 |
| Number of atoms (average B, Å<sup>2</sup>) |  |  |
| Protein |  | 1,285 (65.0) |
| Ligand |  | 902 (55.0) |
| Water |  | 44 (46.5) |
| Ramachandran values |  |  |
| Preferred regions, % |  | 97.7 |
| Allowed regions, % |  | 2.3 |

Values in parentheses are for the highest-resolution shell.
$R_{\text{sym}} = \sum |I_{\text{obs}} - I_{\text{avg}}| / \sum I_{\text{avg}}$ ; $R_{\text{work}} = \sum ||F_{\text{obs}}| - |F_{\text{calc}}|| / \sum |F_{\text{obs}}|$ .
$R_{\text{free}}$ was calculated using 5% of data.

